# Can we use routinely collected hospital and GP data for epidemiological study of common hand conditions? A UK Biobank based validation project

**DOI:** 10.1101/274167

**Authors:** Jennifer C.E. Lane, Christian Schnier, Jane Green, Wee L. Lam, Dominic Furniss, Cathie L.M. Sudlow

## Abstract

**Objective:** Routine health records can be of great value in epidemiological and genetic studies if they are able to reliably identify true disease cases, especially when linked to large cohort studies. Little research has been undertaken into whether coding within UK electronic health records (EHR) is able to accurately identify clinical disease cases of common hand conditions. There is therefore a relative paucity of hand surgical research using EHRs due to concerns that cases cannot be accurately identified.

The aim of this study was to investigate the accuracy of hospital and primary care coding of routine EHRs for carpal tunnel syndrome (CTS) and base of thumb osteoarthritis (BTOA). Self-reported disease state as recorded in UK Biobank, a large prospective cohort study was also investigated.

**Methods:** Code lists for each condition were generated by a team of clinicians, clinical coders and epidemiologists. All patients recruited to UK Biobank in one geographical region (Lothian, Scotland) where linked primary and secondary care coded datasets available were included. A decision-making algorithm was designed to define an administratively-confirmed or a clinically confirmed disease case. Patient electronic medical records (EMRs) were independently interrogated by two clinicians and inter-observer reliability calculated.

**Results:** Of the 17,201 Biobank participants in NHS Lothian, 268 had at least one code for CTS and 82 for BTOA. For CTS, 159 cases were confirmed, 100 cases had insufficient information and 9 cases were refuted. Excluding missing data, the positive predictive value (PPV) for true clinical disease cases was 96% for incident disease (90% for prevalent disease; overall 94%).

For BTOA, 27 cases were confirmed, 46 cases had insufficient information, and 9 cases were refuted. Excluding missing data, PPV for incident disease was 81% (prevalent disease 56%, overall PPV 75%). Interrogation of the disease cases with insufficient information noted a large proportion arising from primary care and self-report coding systems.

Analyzing code combinations revealed that secondary care codes had the highest PPV for CTS and BTOA, emphasizing a more robust evaluation of PPV for patients requiring hospital based care. Overall, inter-observer reliability was good, with agreement in 90% of cases (Cohen’s kappa of 0.79) for clinical disease cases in CTS and agreement of 98%, (kappa 0.96) for BTOA.

**Conclusions:** We have demonstrated that coding within UK Biobank is of sufficient quality to enable use of the resource for epidemiological and genetic research into common hand conditions, and that EMRs can be used for manual validation of UK health coding systems. Further work is needed to consider potential regional and interdisciplinary differences in coding practice, in strategies for dealing with missing data in EHRs, and to validate coding of common hand conditions in primary care.

## Introduction

Musculoskeletal disorders of the hand and wrist are common conditions that present to primary and secondary healthcare services, and cause significant pain and work absence [1]. The two most common conditions in this category are carpal tunnel syndrome (CTS) and basal thumb osteoarthritis (BTOA).

CTS is characterised by pain in the hand and forearm and paraesthesiae of the fingers caused by compression of the median nerve at the wrist. It is the most common compression neuropathy, and estimates of population prevalence range from 0.6% up to 16%. Females are more commonly affected than males, and incidence increases with age [2-4]. A study of trends in surgical treatment noted a significant increase in the number of procedures being undertaken in the UK, with this increasing trend predicted to continue in an aging population [5].

BTOA is a term used to describe osteoarthritis of the trapeziometacarpal joint, that may also be associated with arthritic changes at the scaphotrapezial joint and the scapho-trapezo-trapezoid joint. It is characterised by pain, particularly when using the thumb, and significant reduction in function. The prevalence of base of thumb osteoarthritis has been estimated between 15% and 36% in women and 1.7% and 4% in men, with an annual incidence in the UK of 1.3/1000 in those over 16 years [6-9].

Worldwide, routine health record coding is predominantly designed for remuneration of services, but may also be used in epidemiological research if the coding system can accurately identify disease cases. Routinely coded data has been used in other areas of musculoskeletal research to investigate variations in disease incidence, treatment and outcome [10-12]. Routinely collected health data is of significant research interest when linked to cohort studies. Large cohort studies enable investigation of associations between exposures and incident disease within a population through the augmentation of health records with further information, especially socioeconomic and lifestyle factors. In the UK, linkage of routine health data to large prospective cohort studies such as UK Biobank provides a rich resource with screening for risk factors at recruitment, and additional investigations such as imaging and genetic data to further characterise predisposition to disease [13]. Factors found to be associated with disease through analysis of large prospective cohorts provide substantial evidence to stimulate clinical trials that can confirm causality and improve prevention and treatment.

Large volume cohort data is less readily available for surgical hand conditions than for other musculoskeletal surgical disease groups. National registries such as the national hip fracture database (NHFD), and the national joint registry (NJR) for arthroplasty surgery do not yet exist for hand and wrist conditions. Similarly, the NHS Atlas of Variation, an initiative to reduce inequality in healthcare provision only covers the musculoskeletal treatments of total hip replacement, fractured neck of femur and fragility fracture prevention [14]. The use of routine health records is therefore of increased importance, as large scale epidemiological study of hand and wrist disease is only currently possible in the UK through the linkage of NHS data with prospective cohort studies.

There is currently limited evidence investigating the validity of UK hospital and primary care coded data for common hand conditions such as CTS and BTOA. Published epidemiological studies into these conditions do not include validation of disease coding [4, 9, 15, 16]. Some validation work exploring diagnostic coding in osteoarthritis in general has been undertaken in the Catalan primary care database, SIDAP^Q^, but this has not been done for CTS or specifically for BTOA [10, 17].

Previous validation studies of the reliability of disease coding in UK population databases have found a positive predictive values (PPVs) of between 50-90% and 24-100% respectively, depending on the disease under scrutiny [18, 19]. Other validation studies have used GP questionnaires, GP or hospital records, and, interestingly, lower positive predictive values have been found when validating through the use of medical records [20, 21]. Validation of GP and hospital episode data coding for musculoskeletal disease is less frequently addressed than disease areas such as cancer and mental health disorders [18, 22-24]. Furthermore, Hollowell and Jordan have suggested that there is a greater variability in coding practice for musculoskeletal disease, with an underestimation of disease burden by some databases compared to national statistics [25, 26].

The aim of this study was to investigate the validity of health record coding for two common hand conditions (CTS and BTOA) in the UK Biobank database, as an example of large scale epidemiological resource linking with routine UK health records. These conditions were chosen due to their significant health burden and limited investigation at population level. The conditions also contrast in their manner of presentation to NHS secondary services and the breadth of treatment options available within primary and secondary care. It was therefore considered that there was significant interest in establishing the validity of their coding for future epidemiological and genetic analysis. A secondary aim was to investigate the contribution of two potential sources of coding error: administrative error, and clinical error in the diagnostic process.

## Materials and methods

### Included Codes

A comprehensive set of possible codes for CTS and OA was developed for hospital and primary care data, using the OPCS v4.7 classification of interventions and procedures was used to identify participants who have undergone an intervention for disease in secondary care; the International Classification of Disease (ICD) version 10 was used for the coding of disease in secondary care and the READ coding system (versions 2 and 3) for primary care coding of disease and interventions [27-29].

A list was generated using manual selection of codes, with assistance from a team of expert advisors. For OPCS and ICD coding, advice was sought from Health and Social Care Information Centre (HSCIC) clinical classifications service [30] and local clinical coders working in the NHS. For primary care coding, the QOF framework criteria for general practice was referenced [31]. Relevant codes were also taken from the morbidity definition of hand and wrist site-specific musculoskeletal conditions [32]. Finally, clinicians working in the relevant fields of hand surgery, neurology and rheumatology were consulted, in addition to clinical epidemiologists with experience in validation of health coding systems. The final code lists are given in Supplementary Tables 1-3.

**Table 1.**
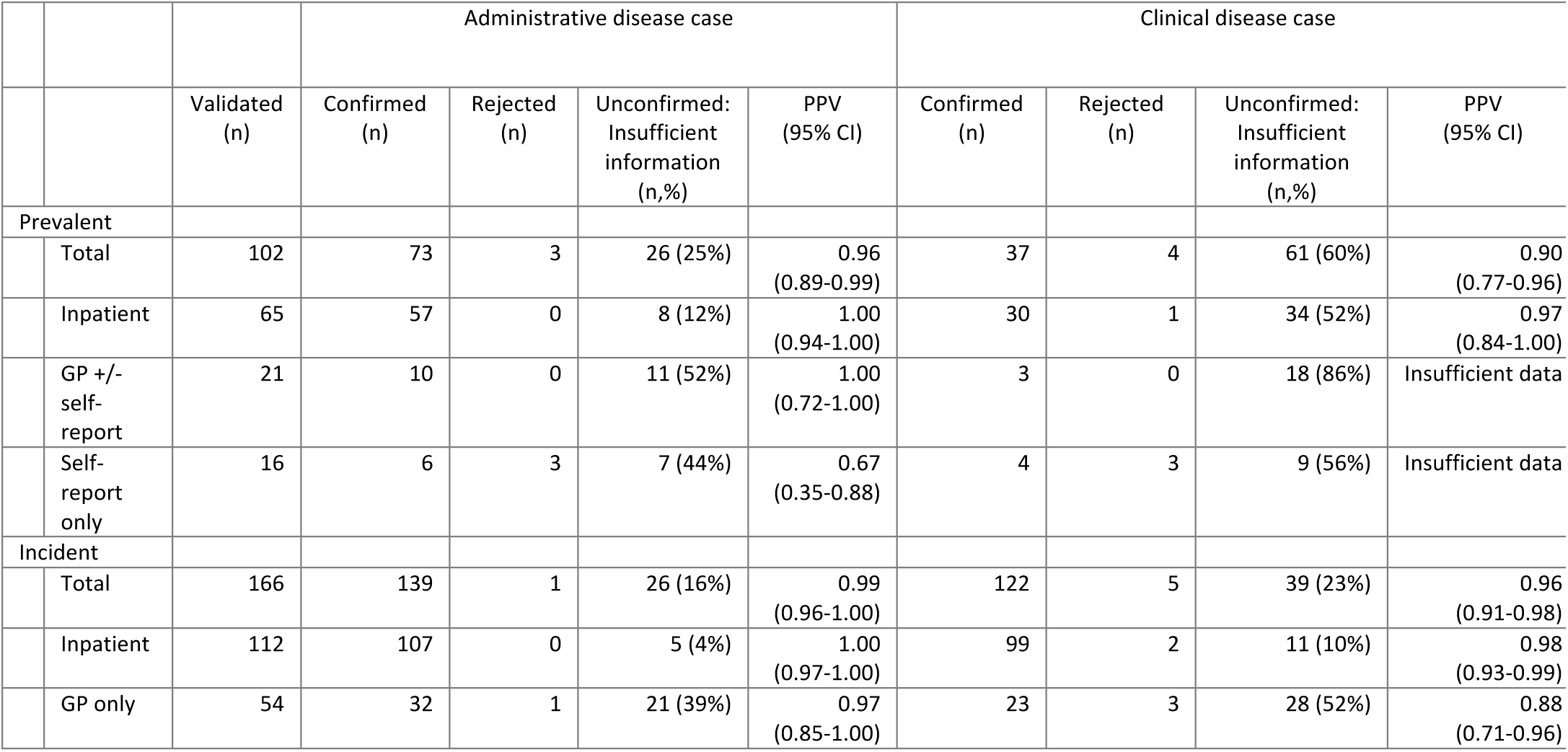
CTS PPV for administrative and disease cases.

### Participants

UK Biobank is a large prospective cohort study of 502,656 people between 40 and 69 years old at Recruitment [13]. Recruitment took place between 2006 and 2010. The UK Biobank database was searched on 14.05.16, at which time, NHS Lothian participants had linked primary care records were linked from January 1990 to April 2013, and secondary care patient records linked from March 1996 to August 2014. We identified all participants in UK Biobank with any code from our list. From this group, all participants recruited in the Lothian region of Scotland were identified. EMRs based in secondary care (TRAK), but containing primary and secondary care data were interrogated. Cases were divided into those first diagnosed after recruitment (’incident‘) and those first diagnosed before recruitment (’prevalent‘). The first disease episode was defined as the earliest date associated with a code in either the primary or secondary care data or in the self-report data. All self-reported disease at recruitment to UK Biobank was considered to indicate prevalent disease, as were any episodes linked to disease documented in the health records prior to the date of recruitment.

### Data collection

Decision-making algorithms were constructed *a priori* to determine the definition of a confirmed or refuted disease case, and an unconfirmed disease case; (where insufficient evidence was found). The algorithm was designed by iterative consensus discussion between epidemiologists and clinicians (Supplementary figures 1-4). The algorithms determined two disease classifications for each participant based on administrative and clinical disease status. These classifications were used to investigate the two main sources of systematic error. The use of an incorrect code was considered to be an ‘administrative error‘; a measurement error associated with the instrument itself, for example a misunderstanding of clinically used terms or a transcription error, causing an inappropriate code to be associated with a patient. A ‘clinical error’ was defined as a patient who was incorrectly identified as a disease case by the clinician through misdiagnosis, leading to a clinical diagnostic misclassification error.

**Figure 1.**
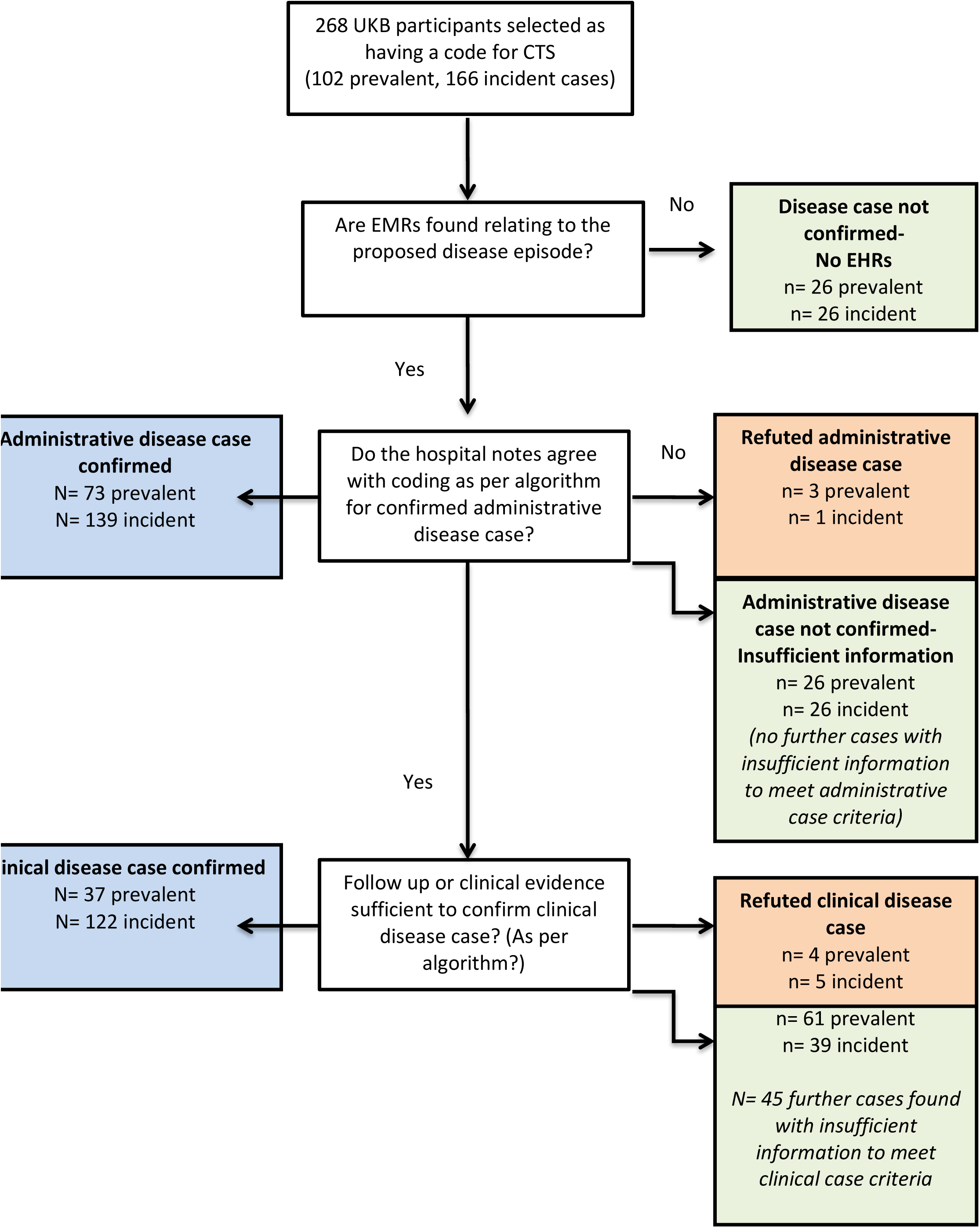
Determining disease case status: CTS.

An orthopaedic surgeon (JL) interrogated the EHR of identified putative cases, with adjudication from a second clinician (CS, a neurologist, for CTS; WL a plastic surgeon specializing in hand surgery for OA.)

### Software

EularAPE software used for basis of proportional Venn diagram representation of results [33].

## Results

Of 17,201 UK Biobank participants from Lothian, 268 people had at least one CTS code and 82 had at least one BTOA code.

Figs 1 and 2 represent how participants were allocated to their disease status for each condition.

**Figure 2.**
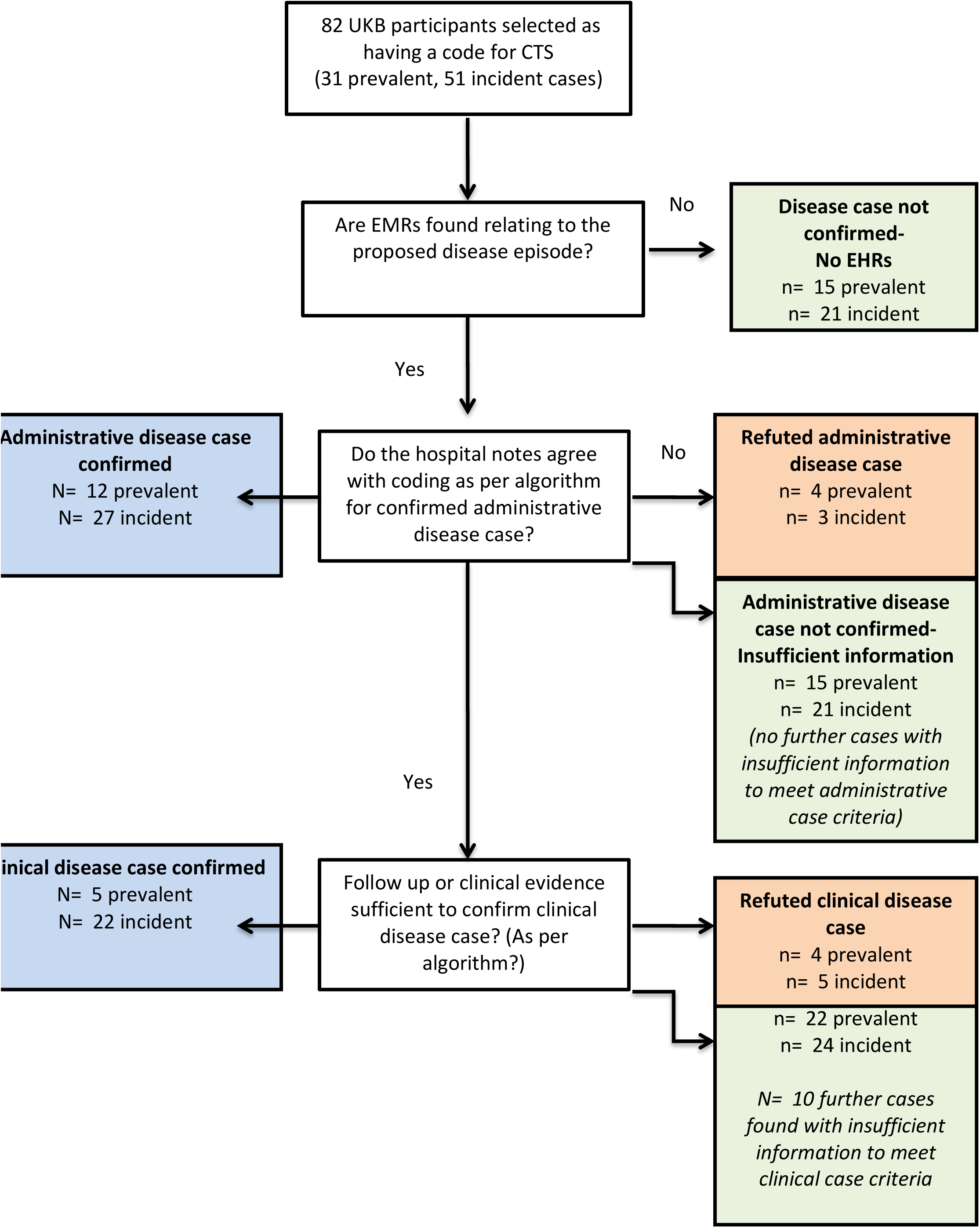
Determining disease case status: BTOA.

Whilst some notes were unavailable electronically, there was also a proportion of participants where EMRs provided inadequate information about the disease episode. The effect of missing EMR information is reflected in the proportion of cases that were defined at each classification level.

**Fig 1. Decision making algorithm**. Results for CTS disease status

**Fig 2. Decision making algorithm**. Results for BTOA disease status

Fig 3 represents the number of cases found between the coding systems for CTS and BTOA, subdivided into prevalent and incident cases. This Venn diagram displays the variation in the burden of disease between primary and secondary care. CTS codes were found to have significant cross over between primary and secondary care; while BTOA cases were predominantly found in primary care. This also emphasizes a potential reason for disparity in the PPV estimates between the diseases, since BTOA was predominantly GP coded, and primary care information was less frequently found in our EMR system.

**Figure 3.**
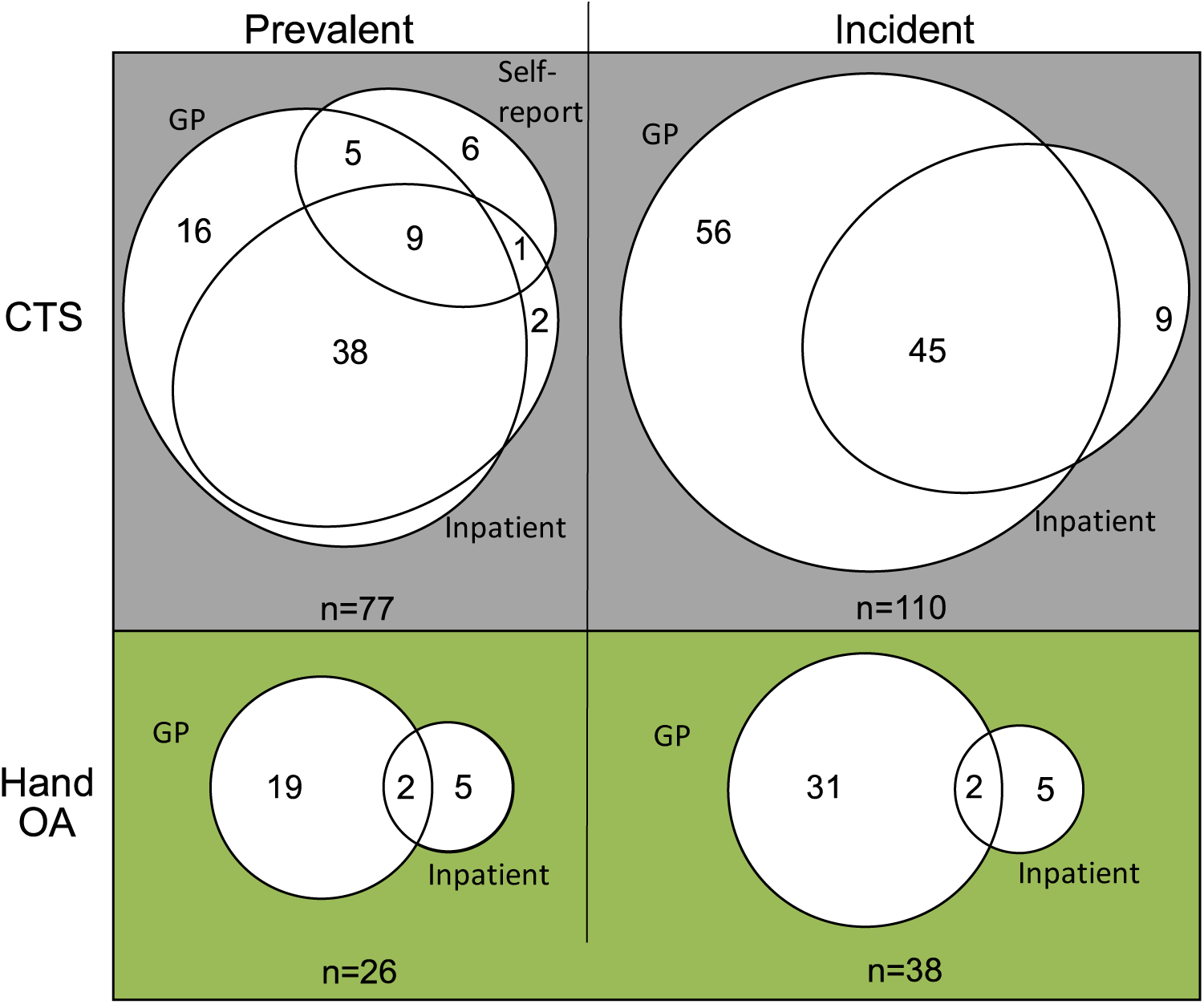
Proportion of cases found within each coding system.

Tables 1 and 2 show the PPV for administrative and clinical disease status for CTS and BTOA respectively.

**Table 2.**
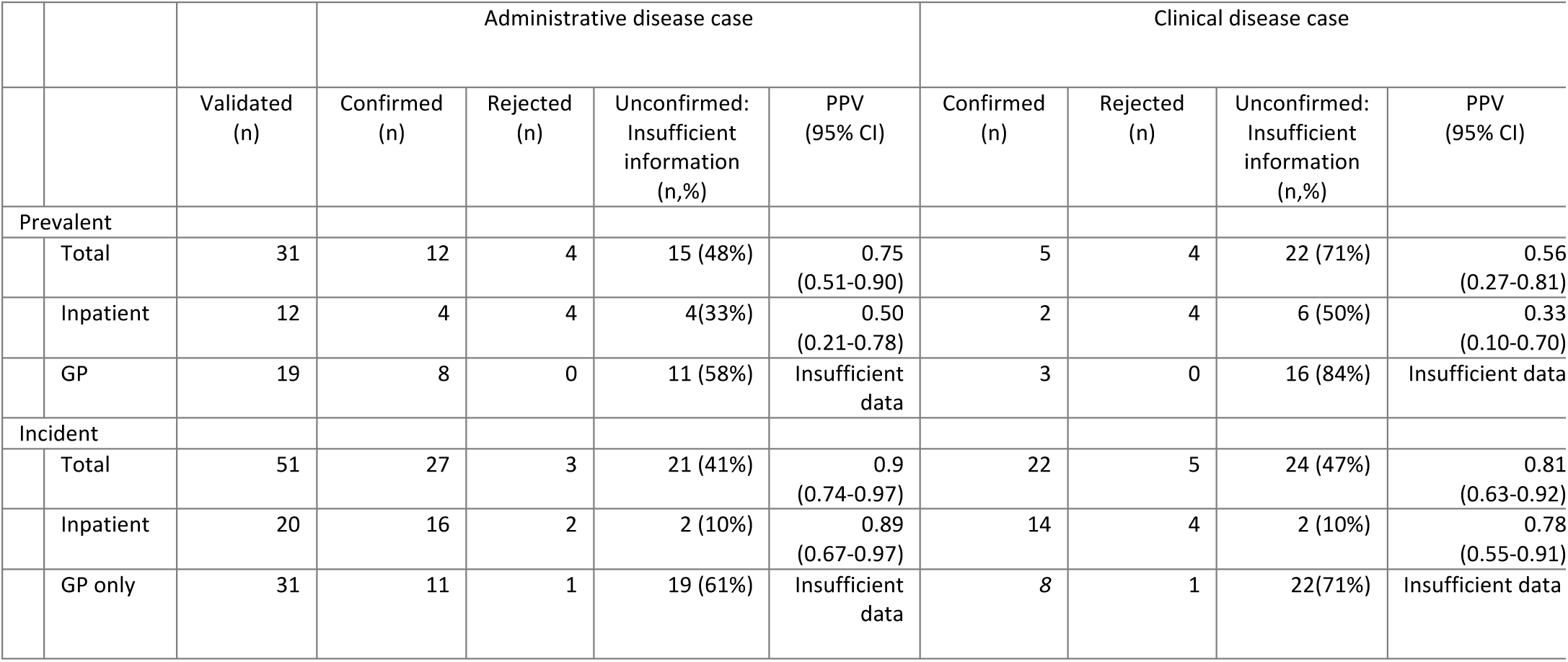
BTOA PPV for administrative and disease cases.

When considering CTS, PPV was highest when using secondary care coding in both administrative and clinical disease cases. This was especially true for incident cases of CTS, where insufficient evidence was found in a low proportion of inpatient coded cases. The proportion of cases with insufficient evidence increased in the primary care coded category, likely to be due to our use of a secondary care based electronic medical record system. Primary care coded cases still have a high PPV for incident disease for CTS, but a large proportion of data was missing in prevalent disease and in self-reported cases. We therefore chose not to evaluate PPV for prevalent primary care and self-reported disease. Considering that the epidemiological value of linking routinely coded healthcare information to cohort studies lies mainly in analyzing associations with incident disease, PPV values for incident cases show that linked healthcare datasets are a robust method of identifying cases of CTS for this purpose.

For BTOA, a greater proportion of unconfirmed cases were seen, leaving a very small number of participants for some coding system combinations. A significant proportion of information was missing for prevalent cases, especially for cases identified through primary care coding. In incident disease, PPV was of a satisfactory, especially in those with inpatient code, indicating that estimates for PPV in patients requiring hospital treatment are more robust. Again, primary care PPVs were not evaluated due to the large amount of missing data.

## Adjudication

Inter-observer reliability assessments for CTS found a Cohen’s kappa statistic of 0.70, with agreement of 94% for administrative disease case confirmation, and a kappa of 0.79 and agreement of 90% for clinical disease case confirmation. Inter-observer reliability assessments for BTOA found a kappa of 0.92, with agreement of 95% for administrative disease case confirmation, and a kappa of 0.96 and agreement of 98% for clinical disease cases. This confirms satisfactory reliability of the clinical expert-led adjudication process.

### Unconfirmed disease status: insufficient evidence found

In order to understand more about the coding process and the potential effect upon positive predictive value, further investigation was undertaken to understand why EHRs were not found for some participants linked to the selected codes.

Unconfirmed cases were predominantly associated with a single primary care or self-report code. This may be due to the time of disease episode being before the introduction of EMRs, or may reflect the reduced level of health care records available from primary care within the regional secondary care-based EMR system. This emphasises the potential for selection bias in this study. Further work is needed to evaluate the accuracy of primary care and self-report data.

As expected, slightly lower final PPVs were found for both CTS and BTOA when using the stricter ‘clinical disease’ definition, in addition to a larger proportion of unconfirmed cases for prevalent disease. This can be attributed to the greater amount of evidence needed to confirm a correct clinical diagnosis rather than a correct administrative code, and the increasing completeness over the time of the EMRs, preventing sufficient detail being available for some prevalent disease episodes.

## Discussion

This study confirms that routinely coded health records are suitable for the genetic and epidemiological studies of carpal tunnel syndrome and base of thumb osteoarthritis. High PPVs are found for both conditions, especially for carpal tunnel syndrome, where if we assume that all unconfirmed cases are not disease cases, and using the precise clinical disease case criteria, the PPV for incident disease was 98% for secondary care coding and 88% for primary care coding. PPVs are lower for BTOA, which in part may be related to the preponderance of primary care coded cases in this group. Primary care data were less well represented within the electronic medical records examined. The lower PPV may also be due to the complex nature of coding for the disease and its associated operative procedures, reflecting the large range of interventions currently used to treat the condition. This contrasts significantly with the more straightforward coding process used for CTS disease and operative intervention.

Within the study we have identified potential sources of error and bias, and attempted to limit them. The difference between the PPVs for the clinical disease and administrative disease cases emphasizes that linkage between database and routine health coding occurs, and that measurement error due to the instrument is not the major source of error. With the health records available in this study, lower PPVs for confirmed clinical disease suggests that the predominant source of error in the use of coded routine health data is that related to misclassification (i.e., error in clinical judgment at the time of diagnosis and intervention).

In order to be useful for epidemiological studies, coded routine health data needs to identify true disease cases. Previous validation studies have concentrated on identifying instrument error. By contrast the use of the more conservative confirmed clinical disease status PPV given here, enables confidence to be taken from associations found when using this data in epidemiological analyses that require a focus on correct clinical disease diagnosis.

Coded healthcare data can also be used to identify disease cases for large-scale genetic research. In this context, the nuance of disease case definition is vital, as variants may predispose to specific clinical sub-phenotypes. It is also important to include only true disease cases in the case group, suggesting that incident cases will give a more tightly defined phenotypic group.

EMRs are increasingly being introduced across the UK and globally, and offer excellent research opportunities, including the conversion of validation into an automated process. Focusing on EMRs in this study does cause some participants to remain unconfirmed in their disease status due to insufficient evidence, but enables us to reflect responsibly about the current limitations of the use of some EMRs as a research resource.

This study is limited to one geographical region, and therefore further work in a different region could increase the number of cases for each condition found, and make the results more generalizable to the whole population. Some coding system combinations had very small numbers associated with them, making it difficult to draw meaningful conclusions.

Manual code selection can be subjective. We attempted to minimize the risk of bias through developing as wide a selection as possible, and using variety of different expert opinions from different relevant clinical specialties to broaden our perspective. The inter-observer reliability was high, but with the difference between expert adjudicators emphasizing the nuance of clinical diagnosis, especially between medical specialties.

## Conclusion

Routine health data from the UK can be used in large-scale epidemiological and genetic research into carpal tunnel syndrome and base of thumb osteoarthritis with confidence. This study highlights the high PPV for both conditions, and emphasizes the utility of linkage of routine data to large prospective cohort studies. Future work is needed to explore primary care coding, to investigate the role of regional variation and to externally validate these results.

## Acknowledgements

The authors would like to acknowledge Lucy Popplewell in the preparation of EMR notes for this study.

The authors would like to acknowledge Keele University’s Prognosis and Consultation Epidemiology Research Group who have given us permission to utilise the morbidity definitions (©2014). The copyright of the morbidity definitions/categorization lists (©2014) used in this publication is owned by Keele University, the development of which was supported by the Primary Care Research Consortium; For access/details relating to the morbidity definitions/categorisation lists (©2014) please go to www.keele.ac.uk/mrr.

**Supporting information**

**S1 Table. ICD code list.** Codes used to identify participants with secondary care disease code for CTS and BTOA.

**S2 Table. OPCS Code list.** Codes used to identify participants with procedure code in secondary care for CTS and BTOA.

**S3 Table. READ code list.** Codes used to identify participants with primary care code for CTS and BTOA.

**S1 Fig. Decision making algorithm: CTS administrative disease case**

**S2 Fig. Decision making algorithm: CTS clinical disease case**

**S3 Fig. Decision making algorithm: BTOA administrative disease case**

**S4 Fig. Decision making algorithm: BTOA clinical disease case**

